# Seasonal variability in temperature, oxygen, and conductivity on the nearshore benthos around McMurdo Sound, Antarctica

**DOI:** 10.64898/2026.09.13.751331

**Authors:** Amy L. Moran, Ming Wei Aaron Toh, Graham T. Lobert

## Abstract

The nearshore marine benthos in McMurdo Sound is one of the best-studied high Antarctic marine ecosystems, yet multiparameter environmental datasets that describe both seasonal and geographic variation in key environmental parameters are lacking. We installed oxygen, temperature, and conductivity loggers at nearshore shallow benthic sites around McMurdo Sound to obtain a high-resolution year-long record of these parameters in 2021 and 2022. Our data showed that benthic organisms experienced a suite of seasonal changes, particularly in temperature and oxygen saturation, that had potential to significantly impact physiology. High summer temperatures were always accompanied by high oxygen saturations, which could provide a metabolic refuge for Antarctic benthic organisms whose oxygen demand increases strongly with temperature. Temperature and oxygen also fluctuated rapidly; on a timeframe of hours to days, these fluctuations sometimes spanned nearly the entire range of seasonal variability. Climate-driven changes to the Southern Ocean are likely to lead to increased seasonal warming, freshening, and potentially deoxygenation, and understanding the extent and rate of change that benthic organisms currently experience will be important to predicting their responses to future conditions.

## Introduction

Understanding natural variation in environmental parameters is a key factor for predicting how organisms may respond to changing environments (1,2). In the Southern Ocean that surrounds Antarctica, even small amounts of change in temperature, salinity, or other factors may have negative implications for organisms that are adapted to millions of years of cold, relatively constant conditions (3,4). In particular, it is a paradigm of Antarctic biology that ectotherms in the Southern Ocean are highly vulnerable to even small amounts of warming (3,5–7), particularly those organisms in the High Antarctic Zone (HAZ) where temperatures are the coldest and least variant (8). Temperature and ocean acidification have received perhaps the most attention as potential threats to Antarctic marine taxa (2,9,10); other aspects of changing oceans like deoxygenation and fluctuating salinity are also likely have far-reaching effects, but have received less attention (11).

Dissolved oxygen is already decreasing in the world’s oceans, with broad biological consequences (12,13). There are comparatively few measurements of oxygen in the Southern Ocean, particularly nearshore (12), however, making it difficult to predict how changes to future conditions will affect the organisms that live there. Warming of the Southern Ocean may create oxygen deficits by increasing metabolic demand, both on an organismal basis and at the regional or ecosystem level (4,14–16), and Antarctic organisms that are adaptively optimized to an environment with a high ratio of oxygen supply to physiological demand may be particularly susceptible to deoxygenation (14,17). Similarly, many subtidal Antarctic organisms are stenothermal; increased melting of sea ice, glaciers and ice shelves is likely to lead to decreases in salinity of nearshore environments, to a degree that will likely be physiologically relevant (18,19). A detailed understanding of the modern environment experienced by near-shore Antarctic marine benthic organisms is therefore essential for both assessing changes, and predicting organismal responses.

Because of its high latitude, seasonal sea ice over, and proximity to the Ross Ice Shelf, McMurdo Sound contains some of the coldest and most thermally stable water in the Southern Ocean and globally (20,21). McMurdo Sound contains many endemic Antarctic taxa (22), and the nearshore benthic communities of the region have been the subject of numerous classic studies on Antarctic ecology (23–29), ectotherm physiology (14,15,30–35), and human impacts (25,36,37). Classically, McMurdo Sound water temperatures were thought to always hover close to the freezing temperature of seawater (Hunt et al., 2003). Though most oceanographic sensor deployments in McMurdo Sound have been relatively short-term due to the logistical difficulty of obtaining data year-round (e.g. Matson et al, 2014), in the past few decades long-term deployments of benthic dataloggers have revealed that there is distinct and biologically-relevant seasonal warming in parts of McMurdo Sound with summer temperature maxima at or around 0 °C (Cheng and Detrich, 2007; Cziko, 2021; Cziko et al., 2014; Hunt et al., 2003). The timing and extent of seasonal warming varies by year and location (Cziko et al., 2014; Kapsenberg et al., 2015; Mahoney et al., 2011), likely due to annual variation in a variety of factors including local and regional patterns of sea ice cover and weather. Salinity measurements are rarer but tend to show a negative correlation with temperature across season (40), probably because summer warming broadly coincides with seasonal sea ice and glacier melt in the region (23).

Like most of the Southern Ocean, oxygen saturation is thought to be high year-round in the water column in McMurdo Sound, with important implications for organismal evolution and physiology (14,15). In the summer, during the seasonal phytoplankton bloom, the surface waters of McMurdo Sound can become hyperoxic (23). In warmer, more studied parts of the ocean, oxygen is often mildly to severely depleted in the benthic boundary layer above marine sediments (41). On the Antarctic benthos, however, organisms are generally considered to have access to high amounts of oxygen throughout the year (42,43), though long-term data records are scarce. Near-bottom and near-shore measurements are also scarce, but one study has shown that under early spring, non-bloom conditions, oxygen was well-mixed throughout the water column at a nearshore site close to McMurdo Station, with typical high (>80% air saturation) levels down to within 1-2 cm of the substrate (44). However, to our knowledge there are no published annual records of seasonal availability of oxygen close to the benthos in McMurdo Sound.

This study investigated the annual patterns of temperature, salinity, and oxygen saturation at several sites around McMurdo Sound, using sensors that were placed ∼3-5 cm above the sediment at shallow (20 - 30 m) nearshore sites to record the conditions experienced by common epifaunal animals like sea urchins, sea spiders, starfish, and invertebrate and vertebrate chordates. Sensor arrays were deployed at different latitudes on both the eastern and western sides of the Sound, which differ in productivity, duration of fast ice cover, temperature and salinity regimes, with corresponding effects on their diversity and benthic ecology (23,26,28,45,46). Typical current patterns in the Sound are dominated by southward flow of Ross Sea water along the eastern side of the Sound (western shoreline of Ross Island) and northward flow of water which is joined by water moving northward out from under the Ross Ice Shelf along the western (continental) side (Fig 1) (23,28,40,45,47) likely leading to persistent differences in the temperature, oxygen, and salinity conditions experienced by the benthic fauna at different localities.

**Fig 1.**
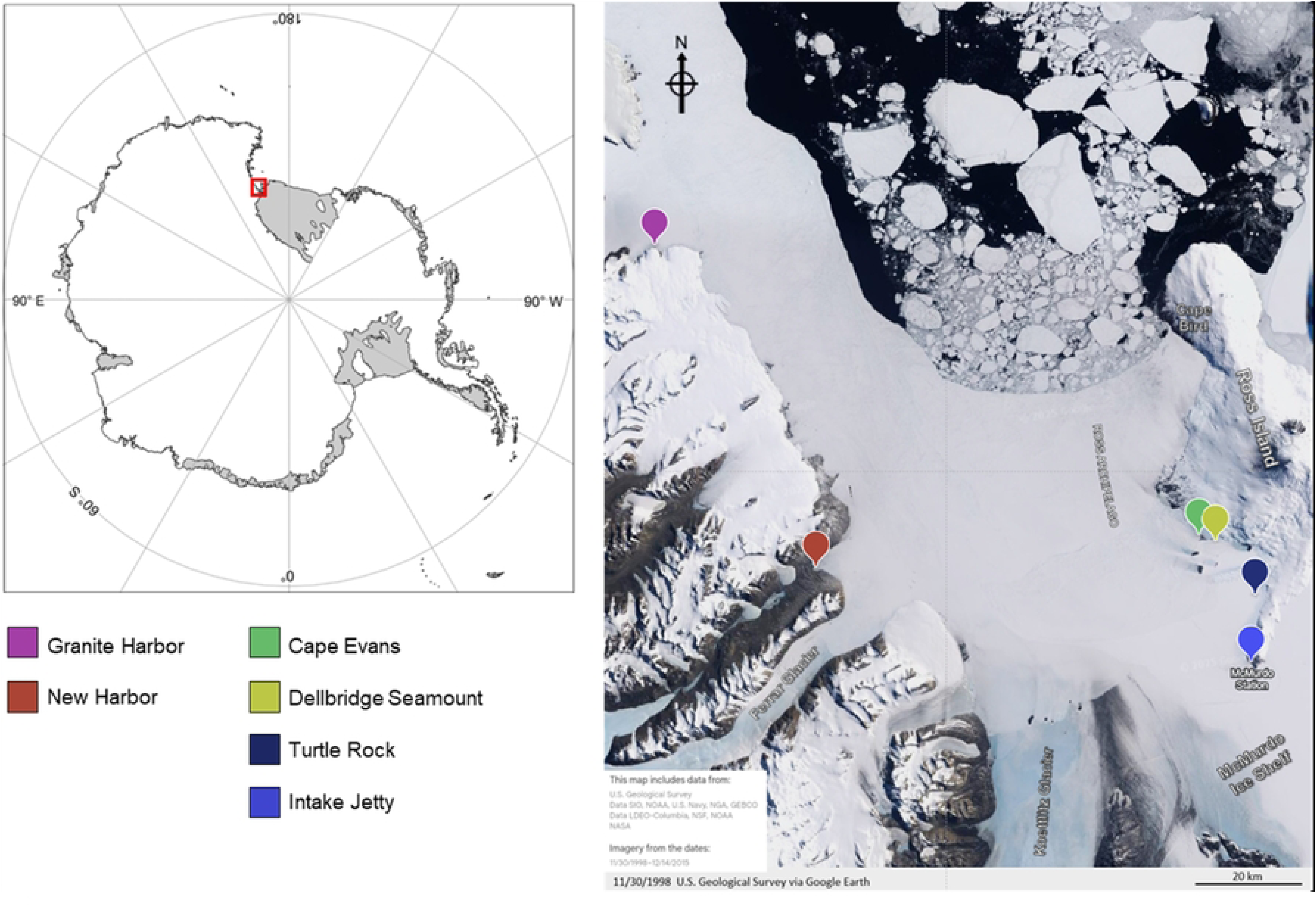
Maps showing the location of McMurdo Sound within Antarctica (left, red box) and logger deployment locations within McMurdo Sound (right). Colored map markers indicate locations of the six sites where loggers were deployed in 2021-2022 (color codes in legend). During deployments, the ice edge was close to Cape Bird on Ross Island (similar to the location in the map image) until late December 2021. The left map image was modified from the Australian Antarctic Division under a Creative Commons Attribution License (CC BY). The right image was modified from Landsat imagery from the United States Geological Survey that are in the US Public Domain.

## Methods

### Datalogger deployment and calibration

Oxygen/temperature and conductivity/temperature sensor/dataloggers (U26-001 and U24-002-C Hobo loggers, Onset, Inc.) were placed by SCUBA divers in the field at several sites on both the east and west sides of McMurdo Sound in the austral spring of 2021 and left in the field until the austral spring of 2022. Sites, logger types, deployment dates, and deployment depths are shown in Fig 1 and Table 1. Sensors were hung from loops attached to a stainless-steel cross-rod installed between two ∼ 1m PVC pipes, one end of each of which was hammered into the substrate (Fig 2). The lengths of the loops were adjusted such that the oxygen sensor heights were approximately 3 cm above the substrate and temperature and conductivity sensors were between 3 and 7 cm from the substrate. To avoid ice formation on sensor surfaces during installation, instruments were kept above 0 °C prior to immersion (Robinson et al., 2020). Loggers took readings every 1 h.

**Fig 2.**
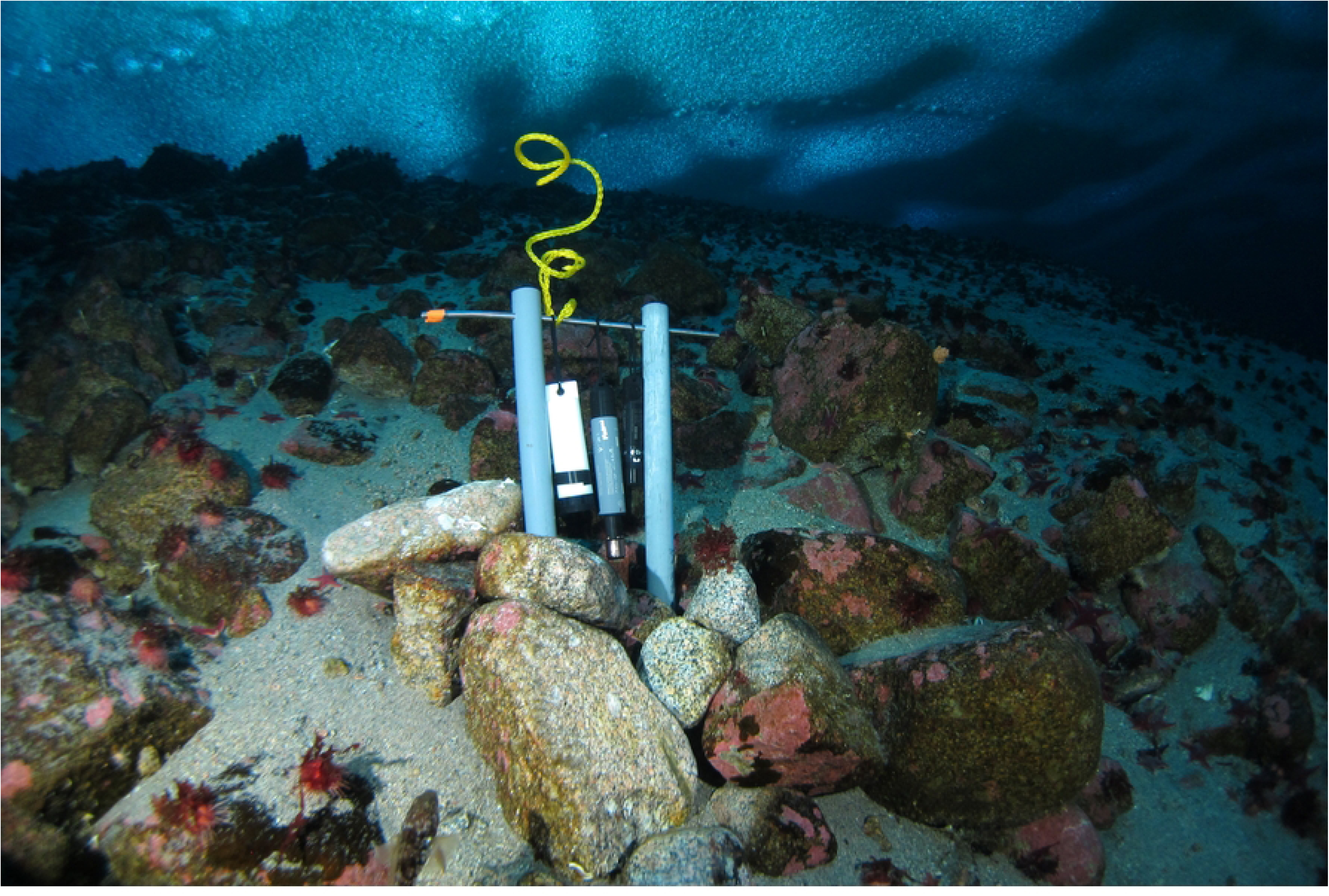
Logger deployment at Granite Harbor. Photo by Rob Robbins.

**Table 1.** Sensor deployment sites, dates, depths, and site characteristics.

| Dive Site name | Coordinates | Start date | End date | Parameters measured | Depth (m) | Site characteristics |
| --- | --- | --- | --- | --- | --- | --- |
| Cape Evans | 77° 38.064' S,<br>166° 24.876' E | 11/02/2021 | 11/13/2022<br>05/17/2022 | Temperature, conductivity<br>Oxygen | 21 | Mix of soft and rocky bottom, invertebrates, macroalgae |
| Dellbridge Seamount | 77° 38.927' S,<br>166° 31.873' E | 11/05/2021 | 11/15/2022 | Temperature | 24 | Mix of soft and rocky bottom, invertebrates, macroalgae |
| Turtle Rock | 77° 44.639' S,<br>166° 46.175' E | 12/03/2021 | 10/7/2022 | Temperature | 27 | Mix of gravel and rocky bottom, invertebrates |
| Intake Jetty | 77° 51.069' S,<br>166° 39.855' E | 10/10/2021 | 10/31/2022<br>09/19/2022 | Temperature, conductivity<br>Oxygen | 21 | Soft bottom, invertebrates |
| New Harbor | 77° 34.268' S<br>163° 30.685' E | 11/23/2021 | 11/24/2022<br>06/22/2022 | Temperature, conductivity<br>Oxygen | 24 | Soft bottom, invertebrates |
| Granite Harbor | 77° 00.715' S<br>162° 51.515' E | 12/07/2021 | 11/29/2022<br>07/07/2022 | Temperature, conductivity<br>Oxygen | 24 | Mix of rocks and gravel, invertebrates, macroalgae |

The temporal range of datasets varied between sites and instruments, for several reasons. First, deployment and collection dates were largely determined by site accessibility, which was in turn determined by operational logistics and sea ice and weather conditions. Second, the U26-001 used optical sensor caps that expire after seven months. Three out of the four sites with deployed oxygen loggers (Cape Evans, New Harbor, Granite Harbor) were completely inaccessible for diving between January 2021 and October 2022, so the sensors could not be replaced and the oxygen records for those sites are seven months or less (the oxygen logger at Cape Evans stopped recording at 6.5 months). At the Intake Jetty, where ice conditions and proximity to McMurdo Station made the site diveable for longer periods, the oxygen logger was collected on 2/15/2022, data were downloaded, the sensor cap was replaced and calibrated, and the logger was redeployed on 2/18/2022. The two datasets from the Intake Jetty logger were merged for analysis.

Prior to deployment, oxygen sensors were air-calibrated according to manufacturer instructions. Oxygen sensors that were still active when collected at the end of deployments (only the two at the Intake Jetty) were immersed in an air-bubbled freshwater ice bath for 24 h while still logging. The measurements taken during this time were assumed to represent true 100% oxygen saturation for each logger and after compensating for salinity differences, the calibration values were used to back-calibrate each downloaded dataset (i.e., if the temperature value in the calibration bath was 99% saturation, 1% was added to each field oxygen estimate from that logger). Temperature was back-calibrated as for oxygen. No calibration was applied to conductivity measurements other than manufacturer calibrations, because due to logistical limitations, high-range calibrations could not be performed at the end of deployments.

### Post-processing and analysis

After post-calibration, data were trimmed to remove readings that were taken in air and the first 10 h of immersed readings, to ensure that we were only using data from instruments that were fully temperature-equilibrated (48). Conductivity readings from the U24-002-C instruments were converted to salinity (g/kg or ppt) using the temperature readings recorded by the same unit and the Conductivity Assistant in Hoboware Pro (v. 3.7.23, Onset, Inc.). Oxygen readings from the U26-001 instruments were converted to percent air saturation using the temperature recorded by the same instrument and the conductivity measured by the U24-002-C sensor in the same array, with the Dissolved Oxygen Assistant in Hoboware Pro. Oxygen levels are reported in units of % air saturation rather than concentration to match the majority of the aquatic organismal and physiological literature (13).

The date and time of the highest (peak) temperature and oxygen measurements were identified from the complete data record from each logger/site by sorting in Excel (Microsoft Office Home and Business 2024, Microsoft, Redmond, WA). Sensor deployment and collection dates differed between sites, so we did not calculate or compare mean temperatures between sites. Likewise, due to the small overall range in temperature (< 2 °C among all sites and dates) and the long duration of deployments, any offset or drift of a sensor’s readings which could not be completely corrected by post-calibration would strongly affect overall means. Similarly, neither peak nor mean salinity values are reported for any site because, due to the overall low variation in salinity and lack of post-calibration, comparisons would be strongly affected by uncorrected instrument drift.

To assess the strength of the relationships between variables within sites, Pearson correlation coefficients for oxygen vs temperature were calculated for each of the four sites that had data records for all three variables (Cape Evans, Intake Jetty, New Harbor, Granite Harbor). To look for potential time lags between changes in oxygen and temperature within site, cross-correlation analyses were performed between oxygen and temperature across the entire temporal sequence at each of the four sites to find the time interval (in h) with the highest correlation.

To assess the time lag between temperature rises on the eastern vs western sides of the Sound, a cross-correlation analysis was conducted on the temperature records for the Intake Jetty (east side) and New Harbor (west side) using only the parts of the two records that overlapped in time (11/23/2021 to 11/21/2022). To address the same question for oxygen levels, a cross-correlation analysis was conducted on the oxygen records for the Jetty and New Harbor sites, again using only the two parts of those records that overlapped in time (Dec. 7 2021 – June 22 2022). These two sites were chosen for the east-west comparison because they were the furthest south in the Sound and experienced the longest periods of fast sea ice cover, so their oceanographic dynamics were most likely to be influenced by large-scale patterns of flow within the Sound rather than local surface warming and bloom conditions. Because of the comparatively small amount of variation in salinity (especially at the Jetty), cross-correlation analyses were not performed between salinity and the other variables.

All correlation and cross-correlation analyses were performed in JMP 18 Pro (v. 18.0.2, JMP Statistical Discovery LLC).

To explore whether the presence or absence of fast ice over the sensors had noticeable effects on oxygen or temperature readings, we looked at daily satellite images of McMurdo Sound acquired from the Visible Infrared Imaging Radiometer Suite from the Suomi National Polar-Orbiting Partnership (VIIRS - SNPP) database, accessed through the NOAA NESDIS STAR Center for Satellite Applications Research Ocean Color Science Team home page (https://www.star.nesdis.noaa.gov/socd/mecb/color/index.php). Three independent observers scrolled through daily images of each site at the highest available resolution (750 m) to identify the first day when there was unquestionably open water (as opposed to fast ice) visible at Cape Evans, Intake Jetty, New Harbor, and Granite Harbor (‘earliest definite’) Because cloud cover often obscured the satellite images, each observer also identified the first day when they thought there was probably open water at each site (‘earliest possible’). Once open water was present, it remained open throughout the rest of the lighted period.

To visualize the relationship between day length (sunrise-sunset) and our measured variables, day length data were obtained from the NOAA Global Monitoring Laboratory using the Sunrise/Sunset and Solar Position Calculator spreadsheet at https://www.gml.noaa.gov/grad/solcalc/calcdetails.html and the GPS coordinates of the most northerly site, Granite Harbor.

## Results

### Temperature

All six sites showed constant, cold winter conditions with temperatures in May – November close to −1.8 °C, followed by a summer rise in temperature starting in December at the sites on the east side of the Sound (Cape Evans, Dellbridge Seamount, Turtle Rock, Jetty) and in January at the sites on the western side (New Harbor, Granite Harbor), returning to winter conditions by May at all sites (Fig 3). At all sites, temperatures were characterized by frequent and often wide fluctuations (compared to the overall seasonal range), sometimes rising or falling by a degree within a day (Supplementary Fig S1). Across the whole season, the highest temperature was recorded at Granite Harbor, followed by Cape Evans, New Harbor, Dellbridge Seamount, the Intake Jetty, and Turtle Rock (Fig 3, Table 2).

**Fig 3.**
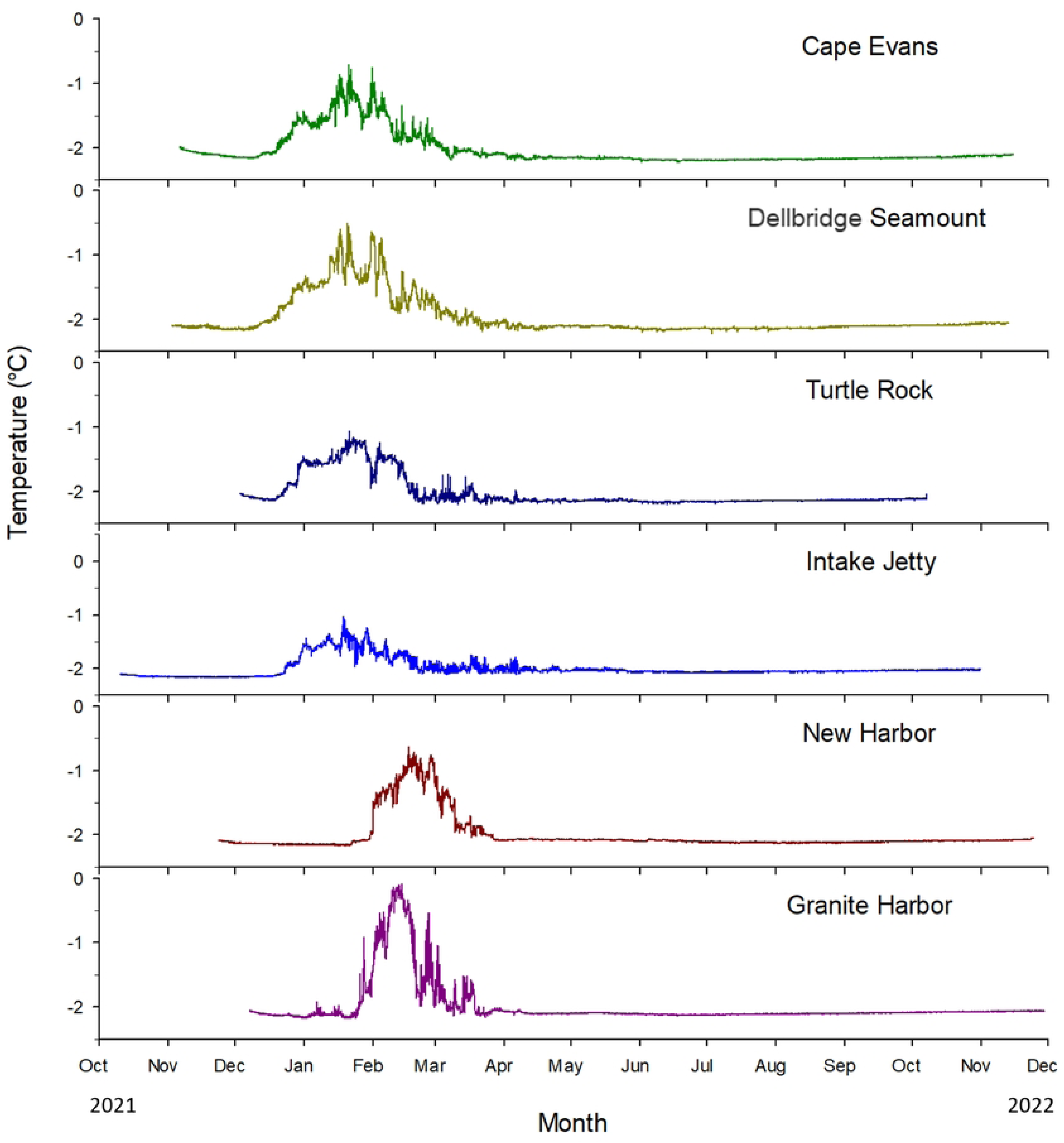
Annual record of temperature from dataloggers placed at six nearshore sites around McMurdo Sound in 2021-2022. Four sites were on the eastern side (Cape Evans, Dellbridge Seamount, Turtle Rock, Intake Jetty) and two on the western side (New Harbor, Granite Harbor). Temperature was recorded every hour. Locations of the sites are shown in Fig 1. Line colors as in site colors from Fig 1. Temperature was recorded every hour. Locations of the sites are shown in Fig 1. Line colors as in site colors from Fig 1.

**Table 2.** Maximum temperatures and dates of occurrence, maximum O_2_ levels and dates of occurrence, Pearson’s correlation coefficients for O_2_ and temperature within sites, and the maximum cross-correlations between O_2_ and temperature with associated lags.

| Site | Max<br>temp<br>(°C) | Date<br>(max temp) | Max<br>O<sub>2</sub><br>(%<br>sat) | Date<br>(max O<sub>2</sub>) | Pears<br>on's r | p | Max<br>cross<br>correla<br>tion | Lag at<br>best fit (h,<br>d) |
| --- | --- | --- | --- | --- | --- | --- | --- | --- |
| Cape Evans | -0.50 | 01/20/2022 | 130.4 | 01/13/2022 | 0.846 | < 0.001 | 0.846 | 0, 0 |
| Dellbridge<br>Seamount | -0.70 | 01/21/2022 | --- | --- | ---- |  | --- | --- |
| Turtle Rock | -1.06 | 01/21/2022 | --- | --- | --- |  | --- | --- |
| Intake Jetty | -1.04 | 01/18/2022 | 135.7 | 01/17/2022 | 0.924 | < 0.001 | 0.936 | -11, 0.5 |
| New Harbor | -0.63 | 02/16/2022 | 121.9 | 02/11/2022 | 0.704 | < 0.001 | 0.754 | -366, 15.3 |
| Granite<br>Harbor | -0.09 | 02/13/2022 | 126.5 | 01/27/2022 | 0.654 | < 0.001 | 0.707 | -128, 5.3 |

On the eastern side of the Sound, temperature began rising at the two northernmost sites, Cape Evans and Dellbridge Seamount, between December 11 and 12, 2021. At the two more southerly sides, Turtle Rock and the Intake Jetty, warming began ∼ eight d later (Fig 3). Temperature first began to rise at the sites on the western side of the Sound more than a month after the eastern side, starting between Jan. 21 and Jan. 22 at both New Harbor and Granite Harbor (Fig 3). The highest cross-correlation between temperatures at the Intake Jetty (the most southerly site on Ross Island) and New Harbor (the most southerly site on the continent) between 11/23/21 and 01/21/22 (these dates encompassed the rise at both sites) was 0.891 and occurred at a lag of 720 h or 30 d (Table 2).

### Oxygen

Oxygen data were recovered from four of the six sites. Dellbridge Seamount was a site of opportunity and no Hobo oxygen logger was installed on the array there. At Turtle Rock, a U24-001 oxygen sensor and logger was initially deployed on the array, but it was missing at the end of the deployment, presumably related to the same disturbance that led to the damage to the array at that site.

At all four remaining sites winter conditions were typified by steady oxygen saturation between 82 and 87% followed by a sharp increase to super-saturated levels (> 100%) beginning in mid-December on the east side of the Sound (Cape Evans, Intake Jetty) and in mid-to-late January on the west side (New Harbor, Granite Harbor) (Fig 4, Table 2). On the western side of the Sound, at New Harbor, oxygen levels showed periodic low spikes from the very beginning of the sampling period, then began to increase on 12/05 and peaked on 01/13. At the Intake Jetty, oxygen remained at winter levels until 12/18 and then abruptly rose to supersaturation over the course of just four days and peaked on Jan. 17. At both sites, oxygen then tapered but remained above winter levels for several months after the main summer pulse, with broader fluctuations evident at Cape Evans. The only site where we were able to identify a complete return to winter levels was the Intake Jetty, where low and constant rates were not reached until around July.

**Fig 4.**
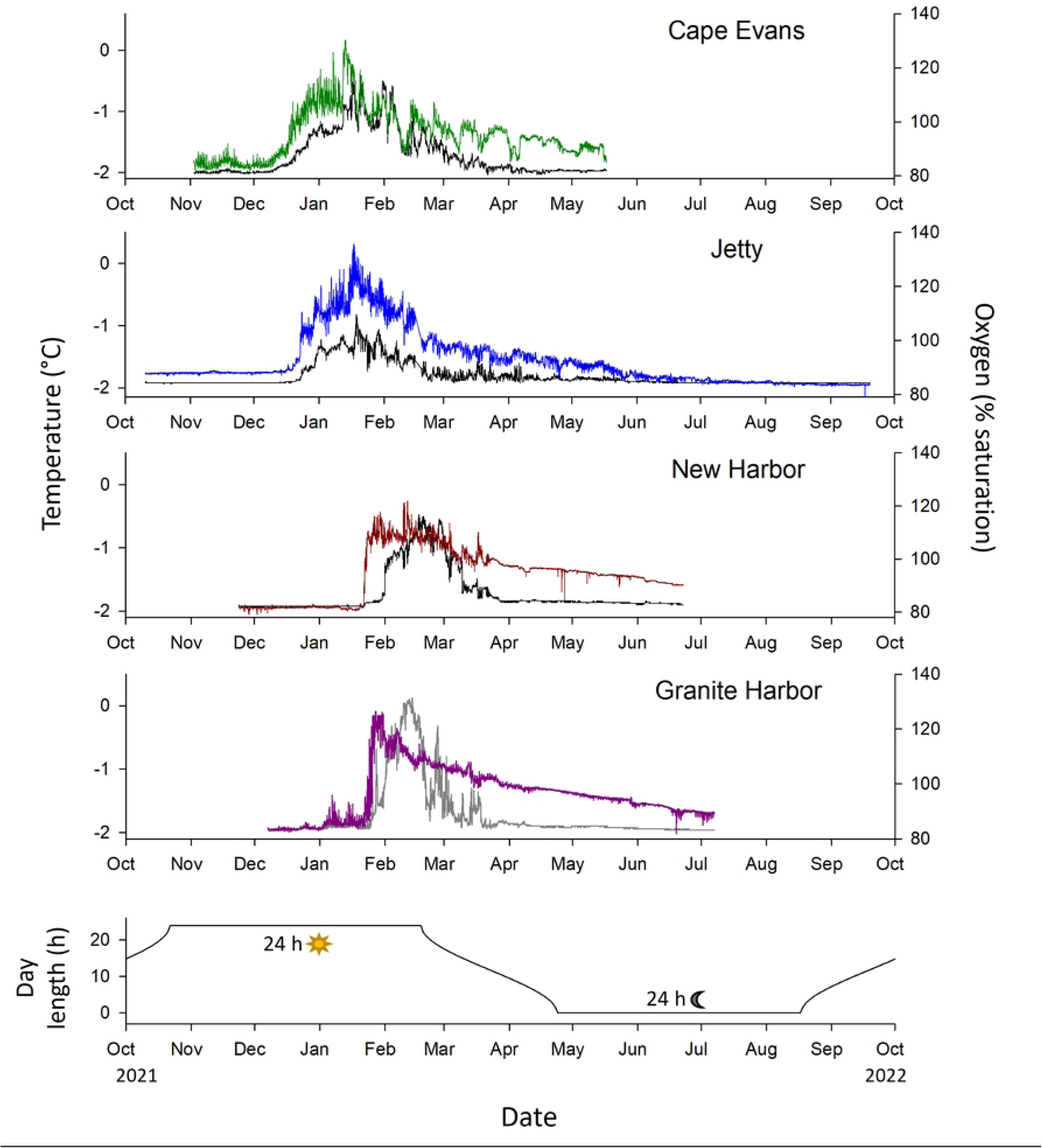
Oxygen, temperature, and day length measured at two sites on the east side of McMurdo Sound (Cape Evans, Intake Jetty) and two on western side (New Harbor, Granite Harbor). Top four graphs: oxygen and temperature measurements from the four sites. Colored lines show oxygen; site colors as in Figs. 1 and 2. Black lines show temperatures measured by the oxygen loggers. Measurements were recorded every hour. Bottom graph: hours with sun above the horizon at Granite Harbor, the most northerly site.

On the western side of the Sound, at the more southerly site (New Harbor), oxygen remained at winter levels until Jan. 21 and then rose to supersaturation over just two d. Peak oxygen levels occurred on Feb. 11. At the more northerly site (Granite Harbor), oxygen was low in December and became more variable in the first three weeks of January. Then, starting on January 22^nd^, oxygen rose from winter levels of ∼83% saturation to the highest levels of the year (∼127%) in just five days. Oxygen levels then showed a steady (though variable) decline until measurements ended in the first week in July, when oxygen saturation still showed some fluctuation and was a few percentage points above the levels in the previous November. Oxygen saturation rose later at the western sites (Fig 4), and the highest cross-correlation between oxygen at the Intake Jetty and at New Harbor between Dec. 7 2021 – June 22 2022 was 0.643, with New Harbor lagging 662 h behind. At all four sites, oxygen rose and peaked at least two months after the onset of 24-hr sunup in late October and remained above pre-bloom levels after the end of April, when the sun was down for 24 h (Fig 4).

### Oxygen and temperature

At all four sites with oxygen measurements, oxygen saturation and temperature were significantly and positively correlated within site (Table 2). Pearson’s correlation coefficients were higher at the two sites on the eastern vs. western side of the Sound. Cross-correlation analyses within sites showed that the best-fit relationships occurred with no lag between oxygen and temperature changes at Cape Evans, and with oxygen lagging 11 h behind temperature at the Intake Jetty, 336 h at NH, and 128 h at Granite Harbor (Table 2).

### Conductivity

Conductivity data were recovered from four sites: Cape Evans, the Intake Jetty, New Harbor, and Granite Harbor. The sensor array at Turtle Rock was knocked over sometime during deployment (probably by seals from the local colony) and the conductivity sensor was partially buried in the gravel substrate upon recovery, so the sensor readings were compromised. The instrument at Dellbridge Seamount was a low-range conductivity meter that we deployed only for temperature readings.

All sites except the Intake Jetty showed comparatively small (1-4 ppt) amounts of freshening beginning in December or January, with salinity returning to winter levels in June or July (Fig 5). On the east side of the Sound, Cape Evans showed a gradual drop in salinity from ∼35 in November to ∼31 ppt in early March, gradually rising back up to ∼34 from mid-April through July. At the Intake Jetty salinity was more constant, showing only a slight freshening from ∼35 to ∼34 ppt over the course of the year and a few very brief (30 minutes-2 h) excursions to ∼30 ppt in June and the beginning of July.

**Fig 5.**
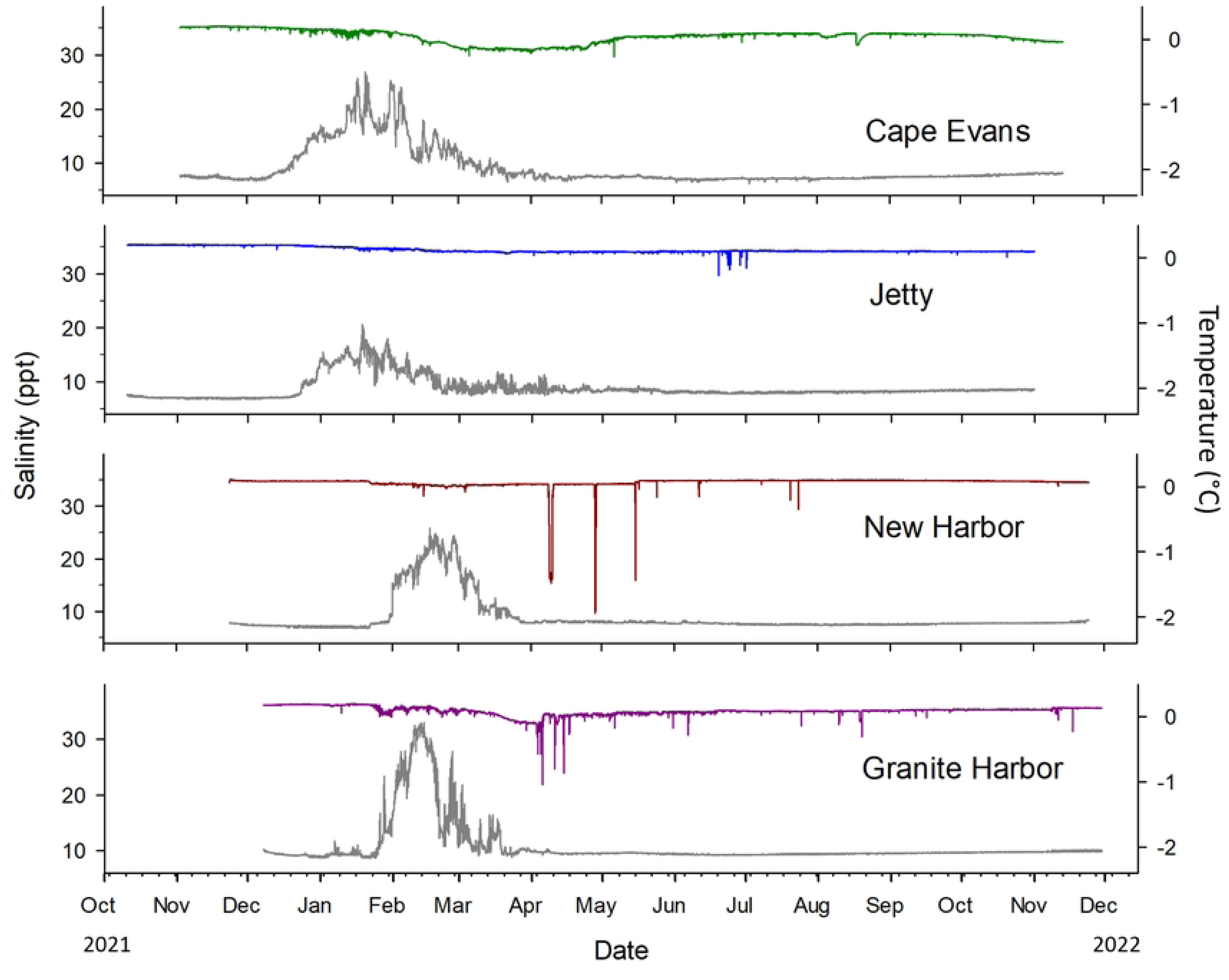
Salinity and temperature data from four sites around McMurdo Sound between Oct. 2021 and Oct. 2022. Colored lines show salinity in ppt; grey lines show temperatures measured by the conductivity loggers. Measurements were recorded every hour. Line colors match site colors in Figs 1, 3, and 4.

On the west side of the Sound, salinity at New Harbor was ∼35 (ppt) in November, December, and early January 2022, followed by a freshening to ∼34 ppt from mid-January through the end of May. This sensor also showed three large and several smaller drops in salinity in April and May. The first and largest of these drops began on April 08 when salinity readings dropped to almost 15 ppt over the course of several hours and remained below 20 ppt for a total of 31 h before rising back up to > 34 ppt by the morning of April 10. The second drop was on April 28, when salinity readings changed from ∼34 to ∼10 ppt in one hour, stayed low for three hours, and then rose back up to ∼34 ppt over the next three hours and remained there. The third and shortest drop was on May 15, when salinity dropped from ∼34 to ∼16 ppt for one hour and immediately rose back up to 33, then 34 ppt. Other than a handful of very short, small decreases, salinity remained close to 34 ppt for the rest of the logging period at New Harbor.

Granite Harbor showed a similar pattern overall to New Harbor, but with a slightly more pronounced drop in salinity between February and May. The logger at Granite Harbor also recorded a handful of excursions to lower salinity around the same time as the New Harbor logger, with salinity dropping briefly to ∼22, ∼24, and again to ∼24 ppt on April 5, 10, and 14 respectively.

### Sea ice extent

Dates when open water appeared at four sites (Cape Evans, the Intake Jetty, New Harbor, and Granite Harbor) are shown in Table 3. In 2021, the fast-ice open-water interface on the eastern side of the Sound was close to Cape Bird, ∼42 km (as the skua flies) to the north of Cape Evans on Ross Island, from mid-September through late December. Open water appeared first at the Cape Evans site in early- to mid-January, and the Intake Jetty site opened in early February. On the west side of the Sound, New Harbor became open water in early-to mid-March and Granite Harbor, early February.

**Table 3.** Dates of earliest likely and earliest definite dates of open water at each site with oxygen data in 2022.

| Site | Earliest possible<sup>a</sup> | Earliest definite<sup>b</sup> |
| --- | --- | --- |
| Cape Evans | January 05 | January 19 |
| Jetty | February 03 | February 11 |
| New Harbor | March 16 | March 17 |
| Granite Harbor | February 04 | February 08 |

\*Satellite images were assessed independently by all three authors; ‘earliest possible’ is the earliest day selected by any of the three authors (variation of 3-5 days).

## Discussion

### Temporal and spatial patterns of oxygen saturation, temperature, and salinity

To our knowledge, this data provides the first annual record of oxygen, temperature, and conductivity recorded simultaneously for shallow benthic habitats at several sites around McMurdo Sound. We identified four main patterns in the data. First, on the east side of the Sound, there was a warm-water pulse that moved from north to south along the west side of Ross Island, starting nearly contemporaneously at Cape Evans and Dellbridge Seamount in mid-December, then appearing at the more southerly sites (Turtle Rock and the Intake Jetty) approximately eight days later. Temperature fluctuated considerably at all sites during the summer pulse, sometimes going up or down by a degree or more within days, especially at the more northerly sites. This warming signal likely represented the influx of warmed surface water from ice-free waters to the north (seasonally warmed Antarctic Surface Water, AASW (49)), advected under the sea ice by the prevailing summer current that brings warmed water as it enters McMurdo Sound from the Ross Sea to the east, curls around Cape Royds, then runs from north to south along the west side off Ross Island (45,50). Consistent with this idea, at both northerly sites temperature peaked around the time that open water appeared at those sites in mid-January. The pattern was very similar at the more southerly sites, except that warming began later and the peak summer temperatures were 0.2-0.5 degrees lower, which could be explained by cooling of AASW as it moved southward and further under the fast ice towards the Ross Ice Shelf. At the Intake Jetty, which has a long history of monitoring and research, the timing, extent, and variability in temperature changes we found were broadly similar to previous annual and multi-year records from the same site (Cheng and Detrich, 2007; Cziko, 2021; Cziko et al., 2014; Hunt et al., 2003).

Second, the comparatively warm summer water mass that appeared on the eastern side of the Sound in December was accompanied by high oxygen saturation, and oxygen and temperature changes were closely coupled. At Cape Evans, oxygen saturation rose gradually from winter levels over the second two weeks of December, it remained close to or above saturation for the next several months, regularly reaching 120% or higher. Because the Cape Evans site was still under ice cover in December, the onset of high oxygen saturation was probably not due to local photosynthetic oxygen production; instead, it probably arose through advection under the sea ice of a large bloom of the diatom *Phaeocystis antarctica* that regularly occurs in open waters north of the Ross Ice Shelf in late October or early November (Arrigo and McClain, 1994; Arrigo and van Dijken, 2004; Rivkin, 1991). This matches previous observations of high concentrations of *Phaeocystis* along with oxygen-saturated or supersaturated conditions along a sampling transect from the ice edge east of Cape Evans to McMurdo Station (54). High variability in temperature and oxygen, accompanied by their close correlation and zero lag at Cape Evans, suggests that there was local mixing of the bloom-derived water mass with colder, lower-oxygen water via eddies, upwelling, or tidally-driven flow at this site. Not much is known about these patterns at the immediate locality of our logger deployments, but all are well known to occur in McMurdo Sound (45,55).

At the Intake Jetty, the most southerly site on the east side of the Sound, oxygen and temperature were again tightly correlated with little lag. Oxygen saturation rose more abruptly at the Intake Jetty than at Cape Evans, increasing by almost 20% over just four days from December 19 to 23. This suggests a more rapid and complete replacement of the winter water with bloom water, which is consistent with decades of diver observations of very rapid decreases in visibility in mid-to-late December at the Intake Jetty site (reviewed in (Hunt et al., 2003). These changes occurred while the Intake Jetty loggers were still under >2 m of fast ice cover, indicating that oxygen was derived from the incoming water mass rather than *in situ* benthic or planktonic photosynthetic oxygen production. While we did not measure oxygen or temperature at more southerly sites closer to the Ross Ice Shelf, if prevailing currents continued to carry the summer bloom water southward and under the ice shelf (as considerable evidence suggests, see Stern et al. 2013), we would predict a pattern of increasingly abrupt arrival of the high-temperature, high-oxygen signal at more southerly sites, along with additional cooling due to longer contact with the sea ice surface and eventually the ice shelf.

The third and most striking pattern was that compared to the eastern side of the Sound, the transition from winter to summer conditions occurred close to a month later on the western side; changes in oxygen and temperature at the two western sites were also much less synchronous than on the east side, with oxygen peaking well before temperature. At New Harbor, the sudden rise in oxygen saturation in late January was almost certainly not driven by local production, because ice cover persisted there until at least early March. If flow patterns in 2021-2022 matched the general observations of cyclonic flow within the Sound (23,45,56), then the rise in oxygen may have signaled the arrival of the same high-oxygen water mass that appeared at the Intake Jetty approximately a month earlier, which had been transported clockwise around the Sound. Interestingly, unlike the east side of the sound, at New Harbor the first sharp rise in oxygen was accompanied by only a small (∼+ 0.05 °C) bump in temperature, suggesting that the water mass experienced cooling during westward transport, possibly due to passing under the Ross Ice Shelf. The rapid rise to close to −1 °C 7-8 days later may have derived from changes to currents, sea ice cover in other parts of the sound, or mixing with warmer water masses from further offshore. At Granite Harbor, though the rise in oxygen occurred around the same time as at New Harbor, it also coincided closely with the appearance of visible open water at that site; thus, high oxygen saturation at Granite Harbor may have been driven both by advection and by local benthic or pelagic photosynthesis. The rise in temperature was also more gradual at Granite Harbor, possibly because its more northerly location is further from the cold-water plume moving northward from under the Ross Ice Shelf (57) and more prone to mixing with warmer water moving westward from northern McMurdo Sound and the Ross Sea.

Like temperature and oxygen, all four sites with conductivity sensors showed some freshening in the summer with later onset at the western sites, and with larger amounts of change at the two more northerly sites. This is consistent with both the cyclonic circulation in the Sound and the earlier transition to open water at the more northerly sites (presumably due at least in part to the melting of sea ice). The sudden, short, and dramatic downward spikes in salinity that we saw at New Harbor and Granite Harbor in April and May seem somewhat unlikely to reflect actual conditions. Very low-salinity water lenses have been reported from under the sea ice at New Harbor and elsewhere in the western Sound in summer (23), but in our data the low-salinity spikes occurred after the melt season; likewise, two separate sensors at each site showed that temperatures remained very steadily at winter temperatures during the low-salinity spikes. While the conductivity sensors performed well during the year-long outplants, these spikes may have resulted from mechanical obstruction of the sensor like transient anchor ice formation or an animal crawling over the sensor surface. We cannot definitively identify the cause, so we chose to leave the data as is for presentation.

### Biological implications

#### Temperature

At all six sites, temperature records indicate that benthic organisms experienced a seasonal warm-temperature pulse that lasted two months or longer and reached temperatures between less than one (Turtle Rock, Jetty) and almost two (Granite Harbor) degrees above typical winter temperatures. It is difficult to say whether these temperature changes would have substantial effects on organismal performance or physiology, for several reasons. First, while Antarctic ectotherms are widely considered to be sensitive to even small amounts of warming (7), many published tests of thermal tolerance of Antarctic animals involved experimental temperatures that were considerably warmer than the highest temperatures we measured in McMurdo Sound (e.g. Flynn and Todgham, 2018; Morley et al., 2024, 2022; Peck et al., 2004; Somero and DeVries, 1967). For those that used similar temperature ranges, results varied; for example, raising temperatures from 0 to +1 °C impaired whole-animal performance of three mollusc species (7) but had negligible effects on the performance of an isopod (60) or a fish (61). Raising temperature from −1 to 0 °C doubled the growth rate of encrusting invertebrates (62) and the metabolic rate of Antarctic krill larvae (63); an increase from −1.5 to +1.5 °C had a similar effect on the metabolic rate of nudibranch embryos (16). Temperature also had a strong accelerating effect on development for sea urchin embryos between −2 and 0 °C (64). It may be that rate processes of Antarctic organisms like growth, metabolism and development are more affected by environmentally-relevant changes in temperature than metrics of whole-organism performance of adults. If so, then the ecological and organismal impacts of current levels of seasonal warming in McMurdo Sound may manifest as changes in reproduction, the development, or organismal energetics, as has already been shown for a subantartic mollusc (65), rather than short-term changes in in performance or individual survival. Within McMurdo Sound, these effects (if present) are likely to be greatest at more northerly sites where temperatures reached higher temperatures, and on the east side of the Sound where the warm-temperature pulse lasted longer.

#### Oxygen

Oxygen saturation on both sides of McMurdo Sound was consistently high (> 80% air saturation) and frequently reached hyperoxic (> 100%) levels during the summer, supporting previous work suggesting that for ectotherms, McMurdo Sound and the HAZ represent a region of high biological oxygen availability (66). When there is a strong effect of temperature on organismal oxygen demand, as for many Antarctic ectotherms (16,67), warming temperatures can cause oxygen deficits if organisms are unable to increase oxygen uptake sufficiently to offset demand (13,68,69). This could impact both whole-animal performance and the developmental rate of Antarctic marine taxa, the majority of which have embryos that develop in benthic egg masses (16,44,70). Unlike adult stages, embryos in egg masses generally rely on diffusion and have few to no mechanisms to increase oxygen uptake when deficits arise(71). Unlike most regions of the world’s oceans, however, our data show that in McMurdo Sound, temperature and oxygen were positively correlated such that the warmest temperatures were almost always associated with hyperoxic conditions. While hyperoxia can have substantial detrimental effects on aquatic organisms (72), it can also rescue development and performance under high-temperature stress (Giomi et al., 2019; Kuchenmüller et al., 2024; McArley et al., 2022; Phillips and Moran, 2015; Wilbur and Moran, 2018; but see Sandrelli et al., 2024). The summer high oxygen saturation in McMurdo Sound may therefore provide a ‘metabolic refuge’ (75) from warm temperature spikes. If so, the combination of high temperature and oxygen may substantially increase developmental rate and biological activity on the benthos in the summer, though these ideas have yet to be tested. If hyperoxia has negative effects, these are most likely to be seen when temperatures have returned to winter levels but oxygen is still high, as was the case for April-June across the Sound.

#### Rate of change

Many organisms adjust to temperature changes in their environment in ways that maximize fitness via acclimation (79), a form of reversible phenotypic plasticity. Antarctic ectotherms are generally considered to be poor acclimators (5,80), which is generally attributed to the low amount of temperature variability that Antarctic ectotherms experience compared to temperate ones (5,6,81). Only a handful of Antarctic taxa have been tested for temperature acclimation. Some of these studies have found no evidence for acclimation (e.g. (81) and others have found that when acclimation occurs, it requires weeks or months (6). It seems unlikely that such slow acclimation would be beneficial in McMurdo Sound, for the obvious reason that by the time an organism had adjusted its physiology to summer temperatures, a return to winter temperatures would be imminent. Another important point relative to acclimation, however, was that in our study both oxygen and temperature fluctuated considerably over quite short time scales; changes that encompassed much of the full seasonal range in both factors often occurred over hours to days. Benthic organisms would therefore experience not just a seasonal increase in temperature and oxygen saturation, but also a rapidly changing temporal landscape in summer where the two factors could change in close tandem (as generally seen at the eastern sites), or at different times (as at the western sites). When organisms cannot reliably predict changes in their environment, theory predicts that phenotypic plasticity is less likely to evolve (82). In addition to the overall stability of Antarctic environments (compared to temperate ones), therefore, the rapid and probably unpredictable nature of changes in temperature and oxygen in nearshore environments may also contribute to the generally low capacity for acclimation of benthic Antarctic ectotherms.

#### Salinity

Because we were unable to fully compensate for conductivity sensor drift with post-deployment calibrations, we did not attempt to compare small differences in salinity between sites or small changes across the full duration of deployment, but rather focused on overall patterns within the salinity record at each site. These patterns suggested that at least in the immediate vicinity of our loggers, particularly at the two southerly sites (New Harbor and Intake Jetty) where sea ice cover lasted until late into the summer, variation in salinity was small and did not reach levels that have been experimentally shown to be physiologically stressful to Antarctic nearshore invertebrates and fishes (18,19). Granite Harbor and Cape Evans (the two more northerly sites) reached lower salinities that could be physiologically significant, when combined with warming; the Antarctic amphipod *Serolis polita* showed no reduction in righting performance when salinity was lowered from 34 to 31 ppt at the control temperature (0 °C), but performance at 31 ppt was substantially reduced as temperatures were raised to 3 °C. In our data, however, while salinity at Cape Evans was close to 31 ppt for almost two months in the late summer, temperatures were always below −1 °C during this interval; similarly, the lowest salinities at Granite Harbor occurred after winter temperatures had returned. Therefore, while experimental tests of the impacts of small changes in salinity on Antarctic taxa are comparatively rare, it seems unlikely that the seasonal salinity changes we saw in McMurdo Sound in 2021-2022 would have major effects on benthic organisms.

Much lower salinities than were seen in our data have been measured in McMurdo Sound, particularly on the west side, where salinity tends to be lower in general and meltwater can form freshwater lenses (23). The logger at New Harbor was deployed in ∼24 m of water close (< 200 m) to the discharge of Wales Stream, where meltwater from the Wales Glacier enters the ocean during the summer. The data from the New Harbor logger suggest that at present, strong physiological effects on benthic organisms from this and similar freshwater discharge points on the west side of the Sound are likely limited to areas that are shallower and/or closer to the immediate vicinity of the discharge. However, as the volume and duration of discharge increase with warming, these impacts will undoubtably increase in both magnitude and area and will expand beyond the currently limited number of sites in the high Antarctic where land runoff reaches the ocean (18,83).

## Conclusions

Nearshore benthic organisms in McMurdo Sound experience annual fluctuations in temperature and oxygen, as well as more rapid fluctuations that are likely driven by mixing of comparatively warm, oxygenated water from summer blooms in the Ross Sea polynya areas with colder winter water and water from under the Ross Ice Shelf. Based on previous physiological studies, these temperature fluctuations were large enough at all sites to impact the rate of important organismal processes like metabolism and development. High temperatures were always accompanied by higher oxygen saturation, which could provide a metabolic refuge from high-temperature stress. High-temperature summer conditions started earlier and lasted longer on the east side of the Sound. An important future direction is to understand how the modern-day high-Antarctic benthic fauna respond to these fluctuations on both seasonal and shorter time scales.

## Acknowledgements

We thank the support staff of the United States Antarctic Program and the McMurdo Station, the USAP diving supervisors and staff for help with sensor deployment and redeployment and dive support, and the staff of the Dive Safety Program at the University of Hawai’i for dive training and support for this project.

## Supporting Information

S1. Hourly temperature recordings over 37 hours from 00:00 January 27 to 13:00 January 28 at Granite Harbor. Data show an increase of > 1 °C over ∼8 h and a subsequent decrease by almost the same amount. The data are a subset of the full annual record from Granite Harbor shown in Fig 3.

## References

1. Gerhard M, Koussoroplis AM, Raatz M, Pansch C, Fey SB, Vajedsamiei J, et al. Environmental variability in aquatic ecosystems: Avenues for future multifactorial experiments. Limnology and Oceanography Letters. 2023;8(2):247–66.

2. Matson P, Washburn L, Martz T, Hofmann G. Abiotic versus Biotic Drivers of Ocean pH Variation under Fast Sea Ice in McMurdo Sound, Antarctica. PloS one. 2014 Sept 15;9:e107239.

3. Convey P, Peck LS. Antarctic environmental change and biological responses. Science Advances. 2019 Nov 27;5(11):eaaz0888.

4. Ingels J, Vanreusel A, Brandt A, Catarino AI, David B, De Ridder C, et al. Possible effects of global environmental changes on Antarctic benthos: a synthesis across five major taxa. Ecol Evol. 2012 Feb;2(2):453–85.

5. Morley SA, Bates AE, Clark MS, Fitzcharles E, Smith R, Stainthorp RE, et al. Testing the resilience, physiological plasticity and mechanisms underlying upper temperature limits of Antarctic marine ectotherms. Biology. 2024 Apr;13(4):224.

6. Peck LS, Morley SA, Richard J, Clark MS. Acclimation and thermal tolerance in Antarctic marine ectotherms. Davies SA, Dow JAT, Lukowiak K, editors. Journal of Experimental Biology. 2014 Jan 1;217(1):16–22.

7. Peck LS, Webb KE, Bailey DM. Extreme sensitivity of biological function to temperature in Antarctic marine species. Functional Ecology. 2004;18(5):625–30.

8. Bilyk KT, DeVries AL. Heat tolerance and its plasticity in Antarctic fishes. Comparative Biochemistry and Physiology Part A: Molecular & Integrative Physiology. 2011 Apr 1;158(4):382–90.

9. Kapsenberg L, Kelley A, Shaw E, Martz T, Hofmann G. Near-shore Antarctic pH variability has implications for the design of ocean acidification experiments. Scientific Reports. 2015 Apr 9;5.

10. Nissen C, Lovenduski NS, Brooks CM, Hoppema M, Timmermann R, Hauck J. Severe 21st-century ocean acidification in Antarctic Marine Protected Areas. Nat Commun. 2024 Jan 4;15(1):259.

11. Brasier MJ, Barnes D, Bax N, Brandt A, Christianson AB, Constable AJ, et al. Responses of Southern Ocean seafloor habitats and communities to global and local drivers of change. Front Mar Sci [Internet]. 2021 May 13 [cited 2025 Mar 30];8. Available from: https://www.frontiersin.org/journals/marine-science/articles/10.3389/fmars.2021.622721/full

12. Hollitzer HAL, Patara L, Terhaar J, Oschlies A. Competing effects of wind and buoyancy forcing on ocean oxygen trends in recent decades. Nat Commun. 2024 Oct 26;15(1):9264.

13. Woods HA, Moran AL, Atkinson D, Audzijonyte A, Berenbrink M, Borges FO, et al. Integrative Approaches to Understanding Organismal Responses to Aquatic Deoxygenation. The Biological Bulletin. 2022 Oct;243(2):85–103.

14. Moran AL, Woods HA. Why might they be giants? Towards an understanding of polar gigantism. Journal of Experimental Biology. 2012 June 15;215(12):1995–2002.

15. Shishido CM, Woods HA, Lane SJ, Toh MWA, Tobalske BW, Moran AL. Polar gigantism and the oxygen–temperature hypothesis: a test of upper thermal limits to body size in Antarctic pycnogonids. Proceedings of the Royal Society B: Biological Sciences. 2019 Apr 10;286(1900):20190124.

16. Woods HA, Moran AL. Temperature–oxygen interactions in Antarctic nudibranch egg masses. Journal of Experimental Biology. 2008 Mar 1;211(5):798–804.

17. Spicer JI, Morley SA. Will giant polar amphipods be first to fare badly in an oxygen-poor ocean? Testing hypotheses linking oxygen to body size. Philosophical Transactions of the Royal Society B: Biological Sciences. 2019 June 17;374(1778):20190034.

18. Barrett NJ, Harper EM, Peck LS. The impact of acute low salinity stress on Antarctic echinoderms. Proc Biol Sci. 2024;291(2031):20241038.

19. Vargas-Chacoff L, Martínez D, Oyarzún-Salazar R, Paschke K, Navarro JM. The osmotic response capacity of the Antarctic fish *Harpagifer antarcticus* is insufficient to cope with projected temperature and salinity under climate change. Journal of Thermal Biology. 2021 Feb 1;96:102835.

20. Hunt BM, Hoefling K, Cheng CHC. Annual warming episodes in seawater temperatures in McMurdo Sound in relationship to endogenous ice in notothenioid fish. Antartic science. 2003 Sept;15(3):333–8.

21. Jacobs SS, Giulivi CF, Dutrieux P. Persistent Ross Sea Freshening from Imbalance West Antarctic Ice Shelf Melting. Journal of Geophysical Research: Oceans. 2022;127(3):e2021JC017808.

22. Brueggeman P. Underwater field guide to Ross Island & McMurdo Sound, Antarctica, volume 5: Arthropoda [Internet]. n.p.; 1998. Available from: https://peterbrueggeman.com/nsf/fguide/index.html

23. Barry JP. Hydrographic patterns in McMurdo Sound, Antarctica and their relationship to local benthic communities. Polar Biol. 1988 May 1;8(5):377–91.

24. Bosch I, Pearse JS. Developmental types of shallow-water asteroids of McMurdo Sound, Antarctica. Mar Biol. 1990 Feb 1;104(1):41–6.

25. Conlan KE, Kim SL, Lenihan HS, Oliver JS. Benthic changes during 10 years of organic enrichment by McMurdo Station, Antarctica. Marine Pollution Bulletin. 2004 July 1;49(1):43–60.

26. Dayton PK. Toward an understanding of community resilicne and the potential effects of enrichments to the benthos at McMurdo Sound, Antarctica. In: Proceedings of the Colloquium on Conservation Problems in Antarctica. Allen Press; 1972. p. 81–96.

27. Dayton PK, Jarrell SC, Kim S, Ed Parnell P, Thrush SF, Hammerstrom K, et al. Benthic responses to an Antarctic regime shift: food particle size and recruitment biology. Ecological Applications. 2019;29(1):e01823.

28. Dayton PK, Oliver JS. Antarctic Soft-Bottom Benthos in Oligotrophic and Eutrophic Environments. Science. 1977 July;197(4298):55–8.

29. Kim S, Hammerstrom K, Dayton P. Epifaunal community response to iceberg-mediated environmental change in McMurdo Sound, Antarctica. Marine Ecology Progress Series. 2019 Mar 21;613:1–14.

30. Cziko PA, DeVries AL, Evans CW, Cheng CHC. Antifreeze protein-induced superheating of ice inside Antarctic notothenioid fishes inhibits melting during summer warming. Proc Natl Acad Sci USA. 2014 Oct 7;111(40):14583–8.

31. Lane SJ, Moran AL, Shishido CM, Tobalske BW, Woods HA. Cuticular gas exchange by Antarctic sea spiders. Journal of Experimental Biology. 2018 Apr 25;221(8):jeb177568.

32. McClintock JB, Pearse JS, Bosch I. Population structure and energetics of the shallow-water antarctic sea star Odontaster validus in contrasting habitats. Mar Biol. 1988 Sept 1;99(2):235–46.

33. Pace DA, Manahan DT. Cost of Protein Synthesis and Energy Allocation During Development of Antarctic Sea Urchin Embryos and Larvae. The Biological Bulletin. 2007 Apr;212(2):115–29.

34. Pearse JS. Slow developing demersal embryos and larvae of the antarctic sea star Odontaster validus. Marine Biology. 1969 June 1;3(2):110–6.

35. Somero GN, DeVries AL. Temperature tolerance of some Antarctic fishes. Science. 1967 Apr 14;156(3772):257–8.

36. Conlan KE, Rau GH, Kvitek RG. δ13C and δ15N shifts in benthic invertebrates exposed to sewage from McMurdo Station, Antarctica. Marine Pollution Bulletin. 2006 Dec 1;52(12):1695–707.

37. Palmer TA, Klein AG, Sweet ST, Montagna PA, Hyde LJ, Sericano J, et al. Long-term changes in contamination and macrobenthic communities adjacent to McMurdo Station, Antarctica. Science of The Total Environment. 2021 Apr 10;764:142798.

38. Cheng CH, Detrich WH. Molecular ecophysiology of Antarctic notothenioid fishes. Phil Trans R Soc B. 2007 Dec 29;362(1488):2215–32.

39. Cziko P. US Antarctic Program Data Center. 2021 [cited 2023 Jan 2]. High-resolution nearshore benthic seawater temperature from around McMurdo Sound, Antarctica (2017-2019). Available from: https://www.usap-dc.org/view/dataset/601420

40. Mahoney AR, Gough AJ, Langhorne PJ, Robinson NJ, Stevens CL, Williams MMJ, et al. The seasonal appearance of ice shelf water in coastal Antarctica and its effect on sea ice growth. Journal of Geophysical Research: Oceans [Internet]. 2011 [cited 2025 Jan 6];116(C11). Available from: https://onlinelibrary.wiley.com/doi/abs/10.1029/2011JC007060

41. Holtappels M, Kuypers MMM, Schlüter M, Brüchert V. Measurement and interpretation of solute concentration gradients in the benthic boundary layer. Limnology and Oceanography: Methods. 2011;9(1):1–13.

42. Macdonald JA, Montgomery JC, Wells RMG. Comparative Physiology of Antarctic Fishes. In: Blaxter JHS, Southward AJ, editors. Advances in Marine Biology [Internet]. Academic Press; 1988 [cited 2025 Jan 6]. p. 321–88. Available from: https://www.sciencedirect.com/science/article/pii/S0065288108600760

43. Wells RMG, Grigg GC, Beard LA, Summers G. Hypoxic Responses in a Fish from a Stable Environment: Blood Oxygen Transport in the Antarctic Fish Pagothenia Borchgrevinki. Journal of Experimental Biology. 1989 Jan 1;141(1):97–111.

44. Moran AL, Toh MWA, Lobert GT, Ely T, Marko PB. Egg masses and larval development of the Antarctic cephalaspidean snail Waegelea antarctica (Cephalaspidea: Antarctophilinidae), with notes on egg masses of the related Antarctophiline alata. Journal of Molluscan Studies. 2021 Sept 1;87(3):eyab027.

45. Barry JP, Dayton PK. Current patterns in McMurdo Sound, Antarctica and their relationship to local biotic communities. Polar Biol. 1988 May 1;8(5):367–76.

46. Dayton PK, Robilliard GA, Paine RT, Dayton LB. Biological Accommodation in the Benthic Community at McMurdo Sound, Antarctica. Ecological Monographs. 1974;44(1):105–28.

47. Littlepage JL. Oceanographic investigations in Mcmurdo Sound, Antarctica. In: Biology of the Antarctic Seas II [Internet]. American Geophysical Union (AGU); 1965 [cited 2024 Dec 17]. p. 1–37. Available from: https://onlinelibrary.wiley.com/doi/abs/10.1029/AR005p0001

48. Robinson NJ, Grant BS, Stevens CL, Stewart CL, Williams MJM. Oceanographic observations in supercooled water: Protocols for mitigation of measurement errors in profiling and moored sampling. Cold Regions Science and Technology. 2020 Feb 1;170:102954.

49. Falco P, Krauzig N, Castagno P, Garzia A, Martellucci R, Cotroneo Y, et al. Winter thermohaline evolution along and below the Ross Ice Shelf. Nat Commun. 2024 Dec 4;15(1):10581.

50. Stern AA, Dinniman MS, Zagorodnov V, Tyler SW, Holland DM. Intrusion of warm surface water beneath the McMurdo Ice Shelf, Antarctica. Journal of Geophysical Research: Oceans. 2013;118(12):7036–48.

51. Arrigo KR, McClain CR. Spring Phytoplankton Production in the Western Ross Sea. Science. 1994;266(5183):261–3.

52. Arrigo KR, van Dijken GL. Annual changes in sea-ice, chlorophyll *a*, and primary production in the Ross Sea, Antarctica. Deep Sea Research Part II: Topical Studies in Oceanography. 2004 Jan 1;51(1):117–38.

53. Rivkin RB. Seasonal Patterns of Planktonic Production in McMurdo Sound, Antarctica1. American Zoologist. 1991 Feb 1;31(1):5–16.

54. Palmisano AC, SooHoo JB, SooHoo SL, Kottmeier ST, Craft LL, Sullivan CW. Photoadaptation in *Phaeocystis pouchetii* advected beneath annual sea ice in McMurdo Sound, Antarctica. J Plankton Res. 1986;8(5):891–906.

55. Lewis EL, Perkin RG. The Winter Oceanography of McMurdo Sound, Antarctica. In: Oceanology of the Antarctic Continental Shelf [Internet]. American Geophysical Union (AGU); 1985 [cited 2025 June 5]. p. 145–65. Available from: https://onlinelibrary.wiley.com/doi/abs/10.1029/AR043p0145

56. Robinson NJ, Williams MJM, Barrett PJ, Pyne AR. Observations of flow and ice-ocean interaction beneath the McMurdo Ice Shelf, Antarctica. Journal of Geophysical Research: Oceans [Internet]. 2010 [cited 2025 June 7];115(C3). Available from: https://onlinelibrary.wiley.com/doi/abs/10.1029/2008JC005255

57. Haas C, Langhorne PJ, Rack W, Leonard GH, Brett GM, Price D, et al. Airborne mapping of the sub-ice platelet layer under fast ice in McMurdo Sound, Antarctica. The Cryosphere. 2021 Jan 19;15(1):247–64.

58. Flynn EE, Todgham AE. Thermal windows and metabolic performance curves in a developing Antarctic fish. J Comp Physiol B. 2018 Mar 1;188(2):271–82.

59. Morley SA, Chu JWF, Peck LS, Bates AE. Temperatures leading to heat escape responses in Antarctic marine ectotherms match acute thermal limits. Front Physiol [Internet]. 2022 Dec 22 [cited 2025 June 9];13. Available from: https://www.frontiersin.org/journals/physiology/articles/10.3389/fphys.2022.1077376/full

60. Janecki T, Kidawa A, Potocka M. The effects of temperature and salinity on vital biological functions of the Antarctic crustacean Serolis polita. Polar Biol. 2010 Aug 1;33(8):1013–20.

61. Sandersfeld T, Davison W, Lamare MD, Knust R, Richter C. Elevated temperature causes metabolic trade-offs at the whole-organism level in the Antarctic fish Trematomus bernacchii. Journal of Experimental Biology. 2015 Aug 1;218(15):2373–81.

62. Ashton GV, Morley SA, Barnes DKA, Clark MS, Peck LS. Warming by 1°C Drives Species and Assemblage Level Responses in Antarctica’s Marine Shallows. Current Biology. 2017 Sept 11;27(17):2698–2705.e3.

63. Quetin LB, Ross RM. Effects of oxygen, temperature and age on the metabolic rate of the embryos and early larval stages of the Antarctic krill *Euphausia superba* Dana. Journal of Experimental Marine Biology and Ecology. 1989 Jan 26;125(1):43–62.

64. Stanwell-Smith D, Peck LS. Temperature and Embryonic Development in Relation to Spawning and Field Occurrence of Larvae of Three Antarctic Echinoderms. The Biological Bulletin. 1998 Feb;194(1):44–52.

65. Reed AJ, Thatje S, Linse K. Shifting baselines in Antarctic ecosystems; ecophysiological response to warming in Lissarca miliaris at Signy Island, Antarctica. PLOS ONE. 2012 Dec 28;7(12):e53477.

66. Chapelle G, Peck LS. Polar gigantism dictated by oxygen availability. Nature. 1999 May;399(6732):114–5.

67. Peck LS, Pörtner HO, Hardewig I. Metabolic Demand, Oxygen Supply, and Critical Temperatures in the Antarctic Bivalve *Laternula elliptica*. Physiological and Biochemical Zoology. 2002 Mar;75(2):123–33.

68. Earhart ML, Blanchard TS, Harman AA, Schulte PM. Hypoxia and High Temperature as Interacting Stressors: Will Plasticity Promote Resilience of Fishes in a Changing World? The Biological Bulletin. 2022 Oct;243(2):149–70.

69. Roman MR, Altieri AH, Breitburg D, Ferrer EM, Gallo ND, Ito Sichi, et al. Reviews and syntheses: Biological indicators of low-oxygen stress in marine water-breathing animals. Biogeosciences. 2024 Nov 14;21(22):4975–5004.

70. Moran AL, Woods HA. Limits to Diffusive O2 Transport: Flow, Form, and Function in Nudibranch Egg Masses from Temperate and Polar Regions. PLOS ONE. 2010 Aug 11;5(8):e12113.

71. Cohen CS, Strathmann RR. Embryos at the Edge of Tolerance: Effects of Environment and Structure of Egg Masses on Supply of Oxygen to Embryos. Biological Bulletin. 1996;190(1):8–15.

72. Welker AF, Moreira DC, Campos ÉG, Hermes-Lima M. Role of redox metabolism for adaptation of aquatic animals to drastic changes in oxygen availability. Comparative Biochemistry and Physiology Part A: Molecular & Integrative Physiology. 2013 Aug 1;165(4):384–404.

73. Giomi F, Barausse A, Duarte CM, Booth J, Agusti S, Saderne V, et al. Oxygen supersaturation protects coastal marine fauna from ocean warming. Science Advances. 2019 Sept 4;5(9):eaax1814.

74. Kuchenmüller LL, Hoots EC, Clark TD. Hyperoxia disproportionally benefits the aerobic performance of large fish at elevated temperature. Journal of Experimental Biology. 2024 Oct 4;227(19):jeb247887.

75. McArley TJ, Morgenroth D, Zena LA, Ekström AT, Sandblom E. Prevalence and mechanisms of environmental hyperoxia-induced thermal tolerance in fishes. Proceedings of the Royal Society B: Biological Sciences. 2022 Aug 17;289(1981):20220840.

76. Phillips N, Moran A. Oxygen production from macrophytes decreases development time in benthic egg masses of a marine gastropod. Hydrobiologia. 2015 Sept 1;757.

77. Wilbur SL, Moran AL. Oxygen-limited performance of the intertidal sea urchin *Colobocentrotus atratus* when submerged. Journal of Experimental Marine Biology and Ecology. 2018 Dec 1;509:16–23.

78. Sandrelli RM, Porter ES, Gamperl AK. Hyperoxia does not improve the acute upper thermal tolerance of a tropical marine fish (Lutjanus apodus). Journal of Experimental Biology. 2024 Nov 7;227(21):jeb247703.

79. Angilletta Jr. MJ. Thermal Adaptation: A Theoretical and Empirical Synthesis [Internet]. Oxford University Press; 2009 [cited 2025 June 13]. Available from: 10.1093/acprof:oso/9780198570875.001.1

80. Peck LS, Morley SA, Clark MS. Poor acclimation capacities in Antarctic marine ectotherms. Mar Biol. 2010 Sept 1;157(9):2051–9.

81. Clark MS, Villota Nieva L, Hoffman JI, Davies AJ, Trivedi UH, Turner F, et al. Lack of long-term acclimation in Antarctic encrusting species suggests vulnerability to warming. Nat Commun. 2019 July 29;10(1):3383.

82. Bitter MC, Wong JM, Dam HG, Donelan SC, Kenkel CD, Komoroske LM, et al. Fluctuating selection and global change: a synthesis and review on disentangling the roles of climate amplitude, predictability and novelty. Proceedings of the Royal Society B: Biological Sciences. 2021 Aug 25;288(1957):20210727.

83. Intergovernmental Panel on Climate Change (IPCC), editor. Changing Ocean, Marine Ecosystems, and Dependent Communities. In: The Ocean and Cryosphere in a Changing Climate: Special Report of the Intergovernmental Panel on Climate Change [Internet]. Cambridge: Cambridge University Press; 2022 [cited 2025 Jan 8]. p. 447–588. Available from: https://www.cambridge.org/core/books/ocean-and-cryosphere-in-a-changing-climate/changing-ocean-marine-ecosystems-and-dependent-communities/EA9AAC25B2A0E030A689E870B42A4E00

